# Understanding the mechanisms of lateral parietal memory modulation in Mild Cognitive Impairment

**DOI:** 10.64898/2026.05.08.723648

**Authors:** Matthew Slayton, Margaret A. McAllister, Emily B. Finch, Kirsten Gillette, Yiru Li, Yuchao Wang, Amanda P. Harris, Jane M. Rothrock, Angel V. Peterchev, Andy Liu, Roberto Cabeza, Simon W. Davis

## Abstract

The application of transcranial magnetic stimulation (TMS) to lateral parietal cortex has shown promise in improving episodic memory in older adults with Mild Cognitive Impairment (MCI). Previous work has suggested that such improvements are achieved by activating hippocampus at a distance with TMS, though this explanation is incomplete. We hypothesized that the mnemonic benefits arise from an additional mechanism: the modulation of semantic representations. Nineteen participants with amnestic MCI received either active intermittent theta-burst stimulation (iTBS) to angular gyrus or control vertex stimulation over three consecutive days while viewing object stimuli and completing relational memory encoding tasks during fMRI, followed by conceptual and perceptual recognition memory tests. We found that active TMS (relative to control TMS) significantly modulated conceptual memory performance. Using Representational Similarity Analysis with semantic embeddings derived from a large language model, we examined how TMS affects neural representations in inferior parietal lobule and hippocampus. We found that TMS enhanced semantic representational strength in inferior parietal lobule and reduced representational strength in hippocampus. Surprisingly, both effects supported successful memory. Neural pattern similarity analyses suggested that reduced hippocampal similarity supported successful memory, perhaps by promoting pattern separation mechanisms. These findings demonstrate that parietal TMS modulates semantic processing in a region-specific manner, by strengthening semantic integration at the stimulation site while promoting representational differentiation in medial temporal regions. This work advances our mechanistic understanding of memory neuromodulation and has implications for the optimization of therapeutic interventions in age-related memory disorders.

**Significance Statement:** Transcranial Magnetic Stimulation (TMS) applied to parietal cortex can improve memory in patients with Mild Cognitive Impairment (MCI), a population at high risk for Alzheimer’s Disease, yet the mechanisms underlying this benefit remain poorly understood. Using fMRI and Representational Similarity Analysis (RSA), we examined how TMS alters the neural representation of semantic stimulus information in parietal cortex and hippocampus during memory encoding. Our results show that TMS selectively modulates semantic representations at the stimulation site and in hippocampus, and that these representational changes predict memory improvement. These findings advance our mechanistic understanding of parietal memory neuromodulation and lay the groundwork for more targeted and effective TMS-based interventions for age-related memory disorders.

## Introduction

Alzheimer’s Disease (AD) is a complex set of brain pathologies that cause decline in memory and cognitive function, affecting over 7.2 million Americans (Gaugler et al., 2025). Mild Cognitive Impairment (MCI) is a condition that has a 30 to 50% chance of converting to Alzheimer’s over 5 to 10 years, and as such is a frequent target for clinical intervention with the goal of slowing progression (Salemme et al., 2025). Over the past decade, there has been increasing interest in modulating episodic memory function in aging by targeting parietal cortex using transcranial magnetic stimulation (TMS), given its network connectivity to regions such as hippocampus (Chen et al., 2022; Hermiller et al., 2019, 2020; Koch et al., 2024; J. X. Wang & Voss, 2015). Moreover, accumulation of amyloid plaques involving parietal cortex is well established (Palmqvist et al., 2017). And yet, fundamental questions remain about the underlying mechanisms through which parietal stimulation affects memory encoding processes. As such, our approach explores the hypothesis that TMS modulates cortical representations of stimulus features, leading to mnemonic benefits.

Successful memory formation depends on the generation and storage of representations of observed experience, a process that is mediated by a widespread cortical network that encodes the visual and semantic features of an experience (Davis et al., 2021). It has been shown that in younger adult populations, TMS applied to angular gyrus (AG) can improve memory performance on item and associative memory tasks (J. X. Wang et al., 2014; J. X. Wang & Voss, 2015). Such studies use a systems level approach in both the selection of a stimulation target (i.e. seeding hippocampal resting connectivity to select the lateral parietal stimulation site) and the primary outcome measure (i.e. memory improvement is correlated with connectivity increases between hippocampus and AG or other Default Mode Network regions) (Hermiller, 2024; Hermiller et al., 2019, 2020). Given that in the earliest stages of AD neuropathological change, Aβ accumulates within the Default Mode Network and impairs connectivity in this region in MCI patients (Palmqvist et al., 2017; Zhang et al., 2025), stimulating a critical node in this network may help restore memory function. While such accounts have provided a mechanism by which parietal TMS affects hippocampal activity, a direct relationship between TMS and stimulus representations has not yet been established.

An emerging literature concerns the capacity of TMS to modulate memory representations. Representational Similarity analysis (RSA) provides the means to address questions of how mental representations are implemented in the brain. TMS applied to dorsolateral prefrontal cortex elicits greater representational strength during both encoding and retrieval in ventral stream regions, an effect linked to both greater hippocampal connectivity and improved item memory (W.-C. Wang et al., 2018). However, evidence that repetitive TMS to parietal cortex modulates semantic representations is lacking, despite the region’s role as a hub for the multimodal integration of sensory information (Humphreys & Tibon, 2023; Ralph et al., 2017; Seghier, 2022; Tibon et al., 2019). In the context of healthy aging, a greater reliance on semantic representational activity can lead to enhanced episodic memory performance (Deng et al., 2021; Naspi et al., 2023), suggesting that neuromodulatory approaches focusing on this region may help to enhance this age-related trend. Moreover, part of the promise of parietal stimulation in age-related memory disorders is that the semantic processing function of AG is preserved across the lifespan, reflecting the general age-related preservation of semantic processing (Cosgrove et al., 2023; Matzen & Benjamin, 2013; Park & Bischof, 2013).

Our goal was to investigate the role of stimulus information in TMS-related memory benefits. MCI participants were scanned during a semantic encoding task following the application of a three-day intermittent theta-burst stimulation (iTBS) to parietal cortex, and semantic representations of object stimuli were assessed using RSA. The cognitive and regional specificity of this effect was tested with conceptual and perceptual memory tests. This constitutes the first application of RSA to understand the effect of TMS on MCI participants, with the goal of understanding how TMS affects the encoding of stimulus information and the subsequent impact on memory, as well as to inform future clinical interventions.

## Methods

### Procedure

This experiment used a between-subjects design with five experimental days including one screening visit which included neuropsychological assessment, one day of functional and structural MRI only and, three consecutive days of TMS followed by MRI. The second visit included a baseline memory task and structural and functional MRI. The results from this task were used to identify the optimal target for stimulation based on subsequent memory effects in lateral parietal cortex. Participants were randomly assigned to the active or control stimulation group. Active participants received intermittent theta-burst stimulation (iTBS) at the lateral parietal cortex site and control participants received vertex iTBS. On the third, fourth, and fifth visits, participants received active or control iTBS after which they immediately completed the encoding phase of a recognition memory task during MRI, followed by the retrieval phase after a short break, repeated for three consecutive visits. All participants underwent functional and structural scans on all four scanning days (**Figure 1**).

**Figure 1.**
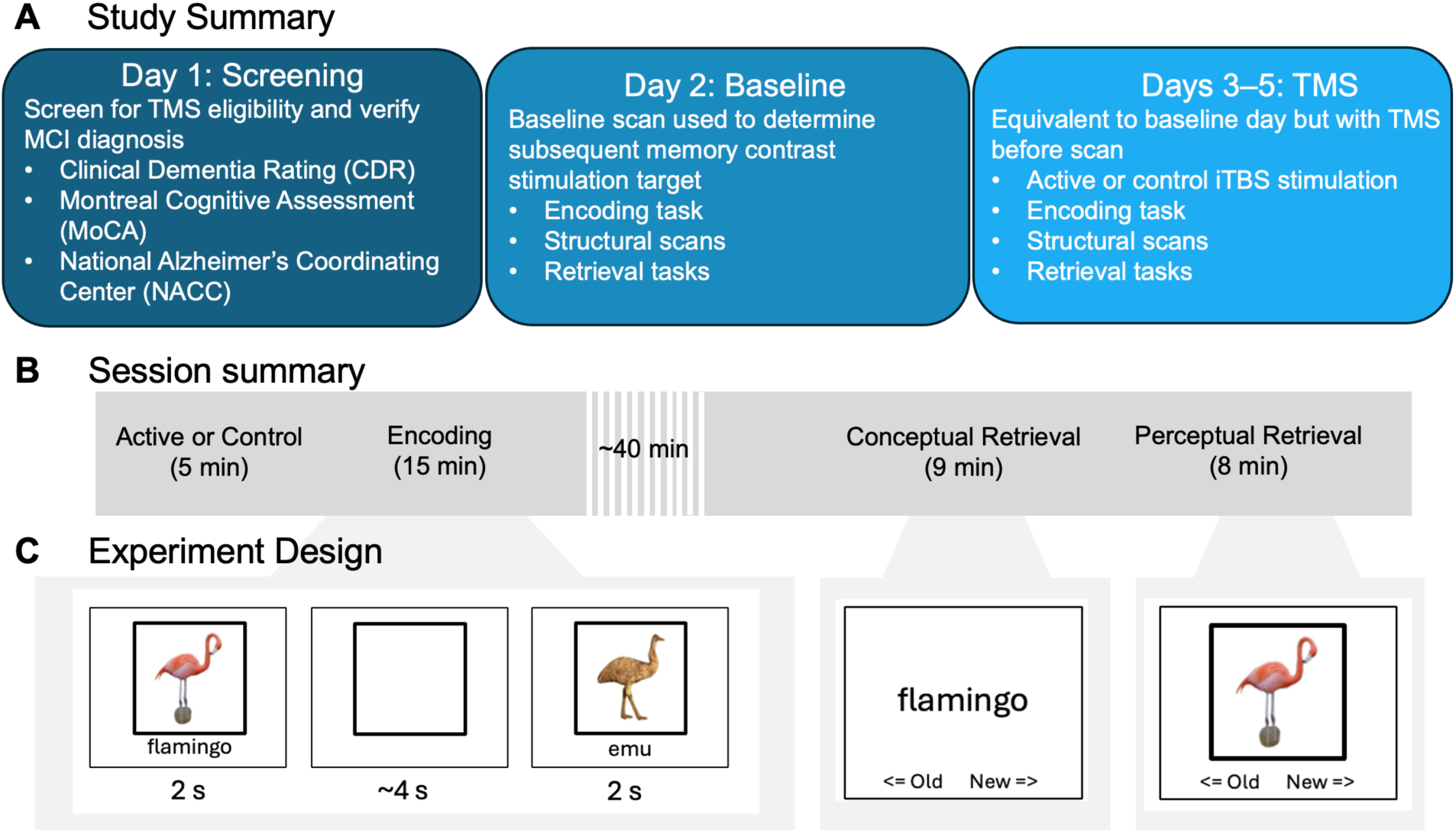
Study Overview. (A) Schematic of overall study design for each treatment day. On screening day participants completed a battery of neuropsychological assessments. On the baseline day, participants completed the memory task in the scanner, and we used a subsequent memory contrast to determine the stimulation target. On the TMS treatment days, participants received parietal (active) or vertex (control) iTBS, followed by three encoding tasks in the scanner. Approximately 40.85 minutes (SD: 11.14 min) later, subjects completed two retrieval tasks for conceptual and perceptual memory. (B) Summary of each experimental session for Days 3–5. Participants received active or control stimulation, performed the encoding task in the scanner, and then performed conceptual and perceptual retrieval tasks. (C) During encoding, subjects were presented with two images with word labels sequentially (separated by a jittered interval of approximately 4 seconds) and were prompted to judge whether the objects were related or unrelated to encourage deep semantic encoding. Retrieval tasks included recognition memory for previously seen or novel objects, presented as either words only (conceptual memory) or images only (perceptual memory). Participants saw objects one at a time and rated them as old or new.

### Participants

The trial was approved by the review board (IRB: Pro00106545) and the local ethics committee of the Duke Medical Center, and all participants provided written informed consent. Participants who received a diagnosis of amnestic Mild Cognitive Impairment were recruited from participating neurologists’ biorepository participant pool, including the Duke Memory Disorders Clinic. All candidate participants were screened for TMS safety in accordance with current guidelines (Chou et al., 2020; Rossi et al., 2021). Any participants who had potential contraindications for TMS were reviewed by the study physician who made the final determination for TMS eligibility. Among our participants there were no contraindications for TMS treatment nor for MRI.

Candidate participants had been given MCI diagnoses prior to recruitment and as part of our screening procedure, we conducted a neuropsychological test battery to establish baseline memory capacity and verify MCI diagnosis. Participants completed the Clinical Dementia Rating (CDR), Montreal Cognitive Assessment (MoCA), and National Alzheimer’s Coordinating Center (NACC) assessments and were classified as having MCI if z scores were 1.5 SD below the mean of the normative data on each metric (Jak et al., 2009). Participants were required to have a MoCA Raw Score of at least 15 to participate in the study.

Data are reported for N = 19 adults (13 females, mean age 73.4 years, standard deviation 4.7 years) who satisfied these criteria and completed the study. One additional participant terminated participation after a recurrent episode of trigeminal neuralgia (McAllister et al., 2023).

### Behavioral Testing

In the relational encoding task, participants viewed word and image pairs one at a time for 2 seconds each, separated by a jittered fixation cross. After the second word-image pair was presented, participants were prompted to make a judgment about whether the preceding objects were related or unrelated, encouraging deep semantic encoding. Participants completed 20 trials (each including two stimuli) in each of three fMRI runs, each of which lasted 5.5 minutes. A mean interval of 40.85 minutes (SD: 11.14 min) separated the end of the encoding task from the beginning of the first retrieval task. The retrieval phase consisted of two recognition memory tasks. First a conceptual memory task, where participants made an object recognition decision with a keypress for individual words presented during encoding that day mixed with novel lures, lasting approximately 9 minutes. Second, a perceptual memory task, where participants made a recognition decision for individual images presented during encoding mixed with novel lures, lasting approximately 8 minutes (**Figure 1**).

### Magnetic Resonance Imaging

Scans were performed on a GE MR 750 3T scanner. We collected functional images using event-related EPI during the encoding task, which was completed in three separate, consecutive runs. Coplanar functional images were acquired using an inverse spiral sequence: 37 axial slices, 64 × 64 matrix, in-plane resolution 4 × 4 mm2, 3.8-mm slice thickness, flip angle = 77, repetition time = 2000 msec, echo time = 31 msec, field of view = 24 mm2. In addition, we acquired full-brain high-resolution T1-weighted structural images with full coverage of the head and neck using a 3D T1-weighted echo-planar sequence (68 slices, 256 × 256 matrix, in-plane resolution 2 × 2 mm2, 1.9-mm slice thickness, repetition time = 12 msec, echo time = 5 mg, field of view = 24 cm).

### Transcranial Magnetic Stimulation

In order to test the differential effects of parietal stimulation on memory for conceptual and perceptual information, behavioral and functional MRI information were collected both before and after treatment with iTBS. In this paradigm, during encoding, participants serially viewed object images and words and judged whether they were related or unrelated. Participants repeated this process for four total days, the latter three of which were consecutive. Behavioral metrics focused on the change in memory performance over the course of the intervention within subjects as well as in comparison to those in the control stimulation condition. We chose a site within lateral parietal cortex defined by the subsequent memory effect, i.e., the difference in activation between remembered and forgotten trials. Vertex (i.e., top of head) stimulation was used for the control condition in a separate group of MCI participants because it plays no active role in memory yet still provides the sounds and physical sensation of stimulation (J. Jung et al., 2016). The subsequent memory approach has some advantages over a resting-state functional connectivity approach (Hermiller et al., 2019, 2020; J. X. Wang & Voss, 2015) in that we have greater specificity about which sites are relevant for memory performance and can therefore infer that activity in these sites contains stimulus-relevant information.

At the start of each TMS visit, participants underwent a urine test to screen for potentially disqualifying medications. They also completed a series of screening questions to gauge sleep and lifestyle factors to assess their mental and physical well-being. Questions were repeated after TMS treatment and at the end of each day to probe for potential side effects of the stimulation.

Resting motor threshold (RMT) was found using frameless stereotaxy (BrainSight: Rogue Research, Montreal) and a figure-eight-shaped coil to identify the minimal power required to induce an index finger twitch. TMS was delivered immediately before the episodic memory encoding task, at 80% RMT using a MagVenture Cool-B65 figure-eight coil positioned over the left angular gyrus (AG) for the active group and over the vertex for control. Stimulation parameters for iTBS comprised 50 Hz triplets at 5 Hz, applied for 2 s, with an inter-train interval of 8 s (c.f. Hermiller et al., 2019). 40 trains were delivered, lasting a total of 4.5 minutes. Parietal stimulation sites include Brainnetome Inferior Parietal Lobule regions: 137: rostrodorsal area 39; 141: caudal area 40; 143: rostroventral area 39. (**Figure 2**).

**Figure 2.**
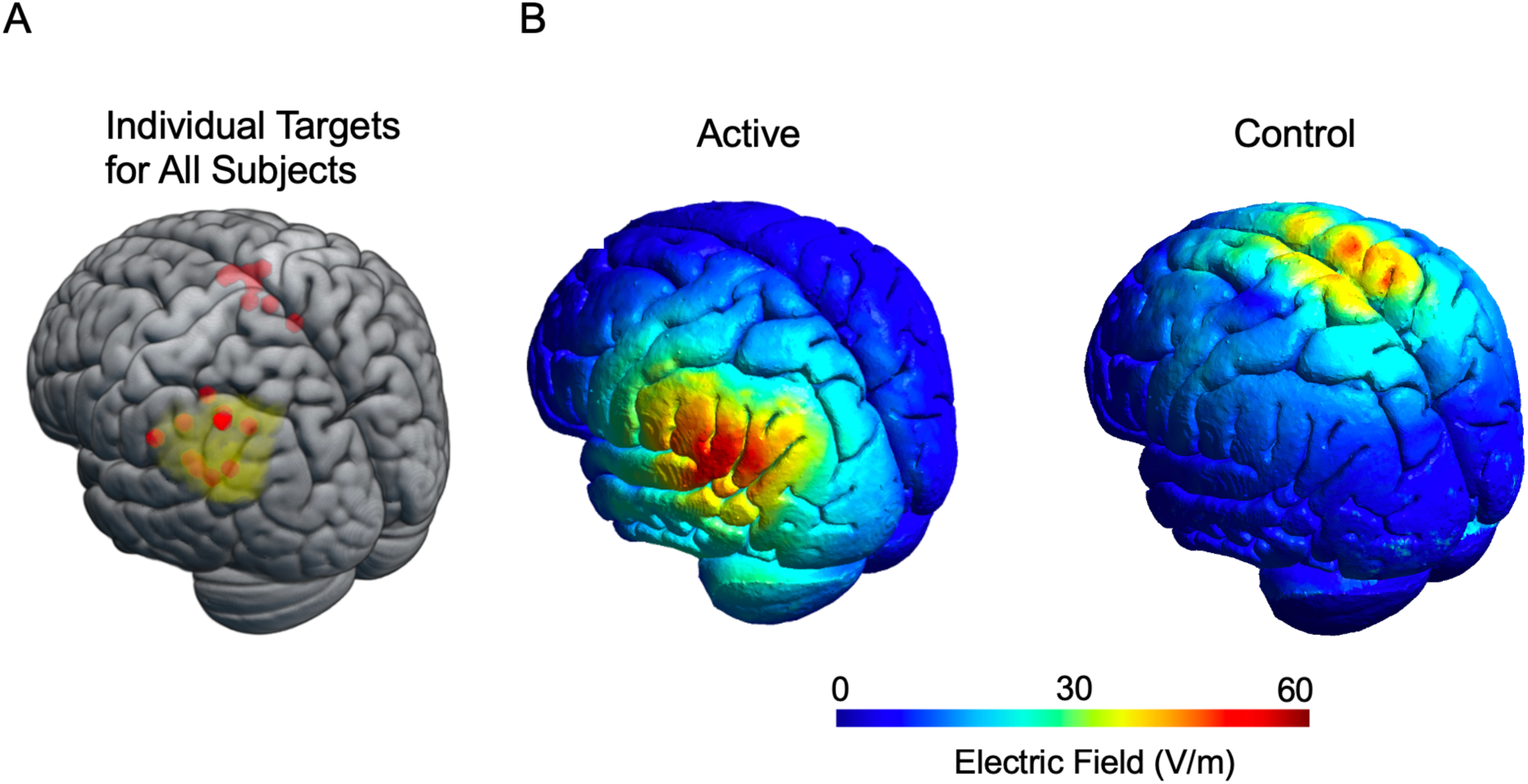
TMS Targeting. (A) Active (n = 10) and control (n = 9) participant target locations, chosen by subsequent memory contrast on conceptual memory task. (B) Electric field (V/m) induced by iTBS at 80% resting motor threshold at either active AG site or control vertex site. After interpolating each participant to MNI space, the E-fields were averaged across the participants in the group.

**Figure 3.**
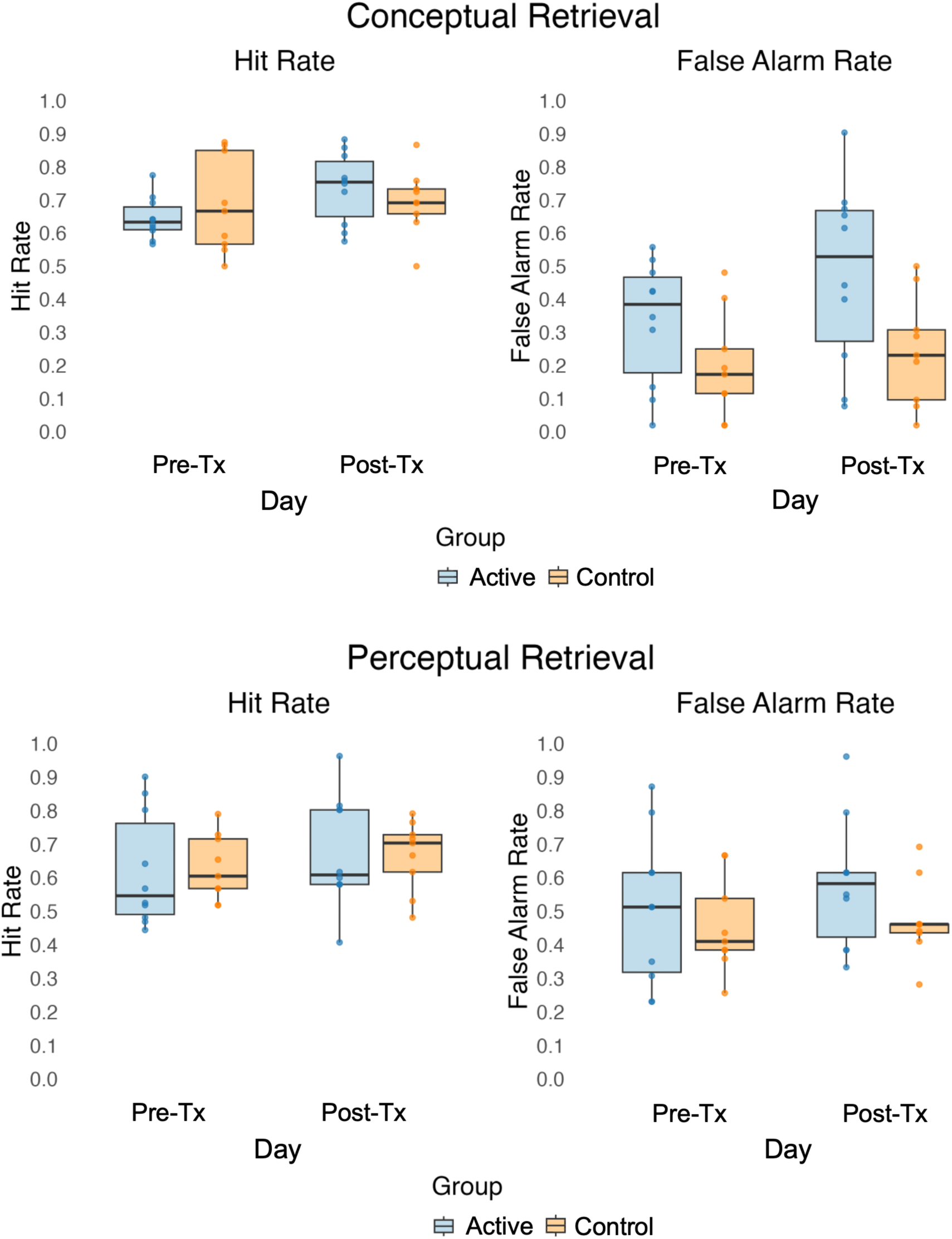
Behavioral Results. Behavioral performance on conceptual and perceptual memory. Subject-wise average performance shown for clarity. (A) Conceptual memory retrieval: hit rate (proportion of hits to hits and misses in response to old items), false alarm rate (proportion of false alarms to false alarms and correct rejections in response to new items), d-prime (difference between normalized hit rate and false alarm rate); (B) Perceptual memory retrieval: hit rate, false, and d-prime.

**Figure 4.**
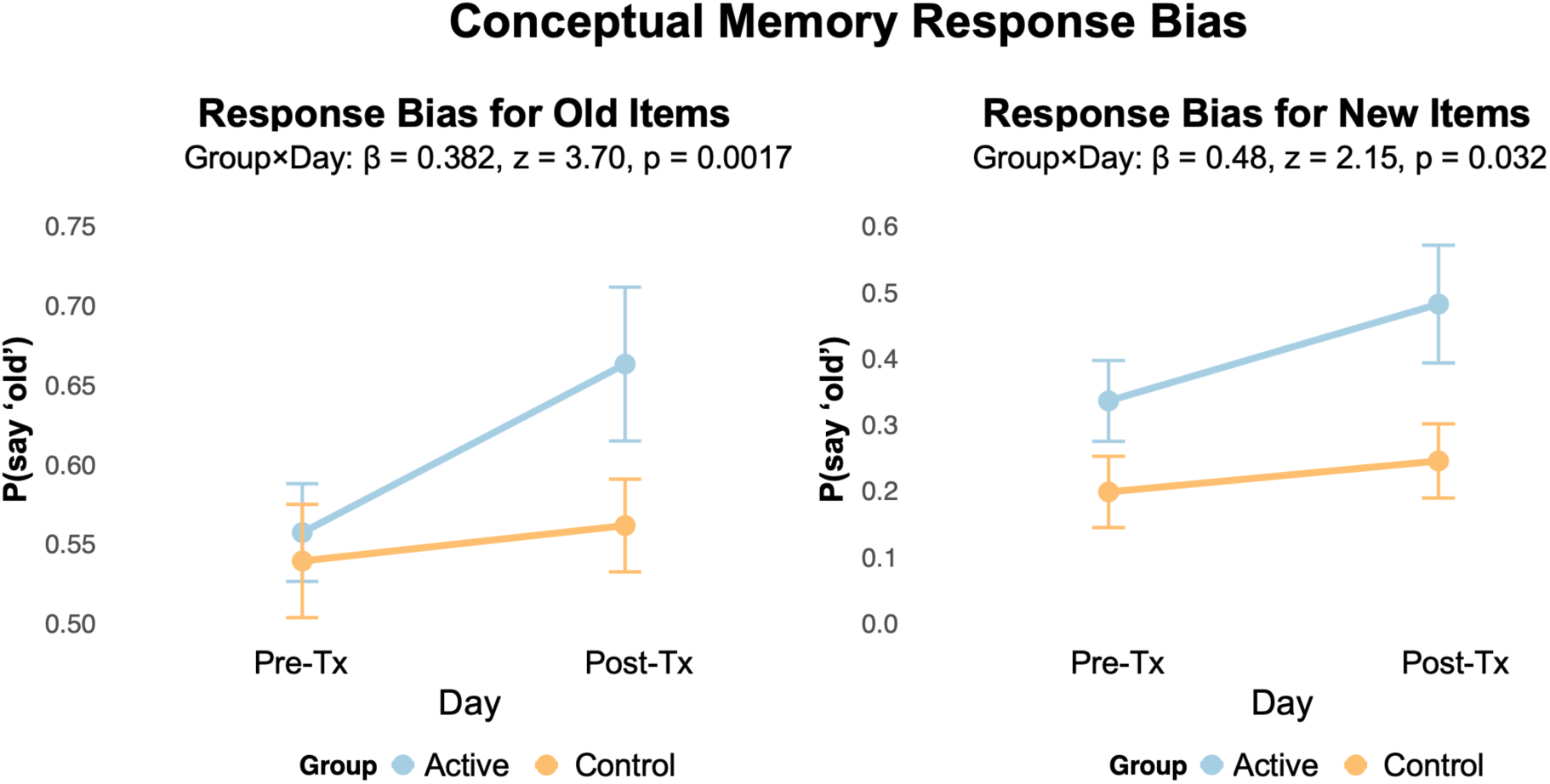
Response Bias. Effect of group and day on response at retrieval for both old and new items.

### Behavioral Analyses

#### Mixed-Effects Models for Hit Rates and False Alarm Rates

Behavioral data were analyzed using mixed-effects linear models implemented in R (version 4.0+) using the lme4 package (Bates et al., 2015) and lmerTest (Kuznetsova et al., 2017) for p-value estimation via Satterthwaite approximations. To investigate group differences in HR and FAR, we used binomial GLMERs: `glmer(Memory ∼ day * Group + (1|subject) + (1|stimulusID), family = binomial)`, where outcome was either hit (for old items) or false alarm (for new items) at the level of trials. Models included fixed effects of treatment group (active vs. control TMS, with control as reference), testing day (1–4), their interaction, and random intercepts for subjects.

#### Baseline-Relative Performance Changes

Baseline performance was calculated as each subject’s mean d’ on day 1 (pre-TMS). We computed two measures of relative change: (1) normalized difference: `(daily d’ – baseline d’)` and (2) percent change: `[(daily d’ – baseline d’) / baseline d’] × 100`. These metrics were analyzed separately to examine both absolute and proportional treatment effects. Effect sizes (Cohen’s d) were calculated for between-group comparisons at each time point using pooled standard deviations to account for unequal sample sizes (n = 10 active, n = 9 control).

### Neuroimaging Analyses

#### Preprocessing

Functional preprocessing and data analysis were performed using fMRIprep version 23.0.1 (Esteban et al., 2019, 2020) and custom Python implementations of the bxh2bids converter, ensuring standardized organizational hierarchy and metadata compliance. Initial preprocessing included removal of the first four volumes (dummy scans) and head motion correction. Data were processed in both subject-native and standardized spaces, with final outputs normalized to MNI152NLin2009cAsym space at both native and 2mm isotropic resolutions. The pipeline included correction for susceptibility distortions, head motion correction, and registration to the anatomical image. ICA-AROMA (Pruim et al., 2015) was applied to identify and remove motion-related independent components. Head motion correction was performed by estimating and regressing out six motion parameters from each functional voxel. Boundary-based registration was disabled due to study-specific requirements. To ensure reproducibility, processing was performed with fixed random seeds based on subject ID numbers, using single-threaded operations where possible. The preprocessing workflow was executed with 16GB of allocated memory and 16 processing threads. All intermediate working files were cleaned upon successful completion of the pipeline. Quality assurance was performed through systematic visual inspection of the preprocessed data, including examination of ICA-AROMA components to confirm appropriate denoising while preserving neurologically relevant signal, and verification of spatial normalization accuracy through visual assessment of anatomical landmark alignment.

#### Univariate Analysis

For univariate analyses, first-level statistical analyses were conducted using FSL’s FEAT (FMRIB’s Software Library, www.fmrib.ox.ac.uk/fsl) and custom MATLAB (The MathWorks) scripts. Individual subject analyses employed a general linear model with local autocorrelation correction (Woolrich et al., 2001). The model included separate regressors for subsequently remembered and subsequently forgotten trials, each modeled as impulse functions convolved with a double-gamma hemodynamic response function. Nuisance regressors included six motion parameters, cerebrospinal fluid signal, white matter signal, framewise displacement, and DVARS. A high-pass temporal filtering cutoff of 100s was applied to remove low-frequency artifacts. For each run, first-level contrasts were computed comparing subsequently remembered versus subsequently forgotten trials. Second-level analyses combined the three runs for each participant using a fixed-effects model. All statistical maps were thresholded at z > 2.7 and registered to MNI152 standard space using FLIRT (FMRIB’s Linear Image Registration Tool). Trial-level activation was computed using the average of beta estimates for each stimulus across voxels within an ROI mask using a general linear model (Worsley & Friston, 1995), as well as a Grey Matter probability mask with a threshold of 0.2. As described previously, a subsequent memory contrast was used to identify candidate targets for TMS.

Next, single-trial beta estimates were obtained using a FLOBS (Flexible Optimal Basis Set with data-driven basis functions) (Woolrich et al., 2004) two-event model. The two regressors were the target object onset (stimulus of interest) and all other onsets from that run (nuisance regressor). This yielded 40 beta estimates per run (20 trials × 2 objects each). All models included standard motion parameters and physiological confounds as nuisance regressors: global signal, white matter, CSF, six motion parameters (three translational, three rotational), root mean square displacement (RMSD), framewise displacement (FD), standardized DVARS, and DVARS. Motion outlier detection was implemented using DVARS > 1.5 and FD > 1mm thresholds to identify high-motion timepoints. All confound regressors were z-score normalized before inclusion in the model.

#### Representational Similarity Analysis

Representational Similarity Analysis (RSA) was conducted using a custom MATLAB pipeline following a trial-specific approach that computed correlations between individual trial patterns rather than comparing full representational similarity matrices (RSM) matrices (Huang et al., 2025). The resulting second-order correlation between rows of the model RSMs and neural RSMs constitutes the *representational strength* for that trial. For each Brainnetome ROI (Fan et al., 2016), neural patterns were extracted from the single-trial beta maps, and similarity was computed between each trial and all other trials from different runs to avoid temporal autocorrelation (in addition to the FILM prewhitening applied during first-level modeling).

These neural similarity patterns were then compared with a model RSM representing semantic properties. In this case, we used layer 6 from Llama (i.e., the seventh layer) 3.2-3b, an autoregressive large language model that includes supervised fine-tuning and reinforcement learning with human feedback. Layer 6 was selected because it is the last of the set of early layers most correlated with item-level semantics (Bogdan, 2025). For visual RSA we used layer 3 of VGG16, the first max pooling layer associated with early visual features such as color (Simonyan & Zisserman, 2015). All similarity values were Fisher Z-transformed and thresholded at z > 2.7 before statistical analysis. To generate the model RSMs, we took word embeddings from Layer 6 and calculated similarity based on Euclidean distance, which we selected after a set of data quality checks meant to choose the most appropriate distance metric (**Supplement S1**).

#### Regions of Interest

For each subject, beta images from each single-trial model were used to estimate activity associated with each object, by averaging all voxel values within a given region of interest (ROI) using the Brainnetome atlas (Fan et al., 2016). Analyses were focused on the left hemisphere because all lateral parietal stimulation was conducted on the left side, consistent with previous work that has targeted angular gyrus with the goal of modulating ipsilateral medial temporal activity (Hermiller, 2024; Hermiller et al., 2019, 2020). We conducted RSA and subsequent memory analyses using two ROIs: (1) Left Inferior Parietal Lobule, a composite of the three Brainnetome ROIs targeted among the 10 active subjects in this study: 137: rostrodorsal area 39 (n = 3), 141: caudal area 40 (n = 5), 143: rostroventral area 39 (n = 2), and (2) Left Hippocampus, including two Brainnetome ROIs 215: rostral hippocampus and 217: caudal hippocampus.

It is well-known that hippocampus supports numerous processes related to the encoding, consolidation, and recollection of episodic memory (Davis et al., 2021; Hainmueller & Bartos, 2020). While the mechanism by which angular gyrus supports memory remains an active area of debate, several studies applying TMS to angular gyrus have found that such intervention can affect memory success, confidence, and vividness (Hower et al., 2014; Richter et al., 2016; Simons et al., 2010; Tibon et al., 2019; Yazar et al., 2012, 2014), which supports the model that AG acts as a hub for the multimodal integration of sensory features into memory (Shimamura, 2011).

##### Representational Strength and Memory Performance

To examine the relationship between neural representations and memory outcomes, we implemented trial-level mixed-effects models analyzing how representational similarity (RSA) values in each of our ROIs predicted memory performance. We modeled both conceptual and perceptual subsequent memory performance as binary outcomes using GLMERs: `glmer(Memory ∼ RSA * day * Group + (1|subject) + (1|stimulusID)`, family = binomial). Memory outcomes were coded as –0.5 (forgotten) or 0.5 (remembered). Models included three-way interactions between RSA strength, treatment day, and group (active vs. control TMS), as well as two-way interactions between RSA and group. Random effects accounted for repeated measures within subjects and items. We then conducted additional, complementary analyses to analyze effects at all levels of our experimental factors. These included `Group:Day` effects for subsequently remembered trials, subsequently forgotten trials, and all trials regardless of memory, as well as `Group:Memory` effects for day 1 (pre-treatment baseline), day 4 (post-treatment). We then investigated group effects at all nine combinations of three levels of memory and day. This multi-level approach allowed us to characterize how TMS effects on neural representations varied as a function of both memory outcome and treatment timing without relying only on two-way interactions that are estimated at the reference level of the third. We decided to focus on a two-way interaction model for subsequently remembered trials rather than a three-way interaction with memory in order to isolate the relationship between Group, Day, and representational strength, rather than investigate the `Group:Day` interaction slope and representational strength at different levels of memory, which would be given by a `Group:Day:Memory` model. For comparison, three-way interaction results are included in **Supplement 3**.

#### Recognition Response Models

To investigate the relationship between response during encoding (where participants make relatedness judgments about the presented image pairs) and the *a priori* similarity of the items, treatment group, and treatment day, we used the following mixed-effects model: `response ∼ day * Group + (1|subject).` We then proceeded to add the neural-behavioral relationships to these models with the addition of ROI-wise RSA: `RSA ∼ response * day * Group + (1|subject)`.

Finally, we investigated the impact of group and day on response at encoding for both old and new items: `response_at_retrieval ∼ Group * day` to see if TMS treatment affected recognition decisions at retrieval. The inclusion of new items makes this model different from the memory success model which by definition uses old items only. We replicated the same approach to handling potential interpretive issues with higher-level interactions as we did in the memory models to estimate the effect of group as well as the effect of group and day interactions at different levels of response at encoding.

#### Multiple Comparisons Correction

P-values were adjusted for multiple comparisons using the Bonferroni-Hochberg (BH) method for each pair of tests (for the two ROIs) (α = 0.05/2 = 0.025). Confidence intervals for fixed effects were computed using the Wald method with degrees of freedom estimated using the Satterthwaite approximation for linear models and z-distributions for generalized linear models.

#### Neural Pattern Similarity Analysis

To investigate whether pattern separation mechanisms support subsequent memory, we computed trial-level neural pattern similarity (NPS) metrics for encoding trials and tested their relationship with later memory performance. We used the same single-trial beta estimates from previous analyses and focused on our two regions of interest: inferior parietal lobule and hippocampus. For both ROIs, we extracted voxel activation patterns across all trials within a session. We then computed NPS, which was a Fisher z-transformed correlation between an item’s pattern and all items from different runs within the same session. This metric is meant to capture general neural pattern distinctiveness, controlling for run-specific effects. We also extracted semantic similarity values for each pair from language model-based embeddings (i.e. the same used to generate the model RSMs for RSA).

#### NPS Statistical Analysis

We tested whether NPS predicted subsequent memory using mixed-effects logistic regression, separately for each ROI and NPS metric. Random intercepts were included for subject and stimulus ID to account for individual differences and item-specific effects. Model 1 tested global similarity: `memory ∼ NPS * Group * Day + (1|subject) + (1|stimID)`, and Model 2 tested semantic moderation: `memory ∼ NPS * pair_semantic_similarity * Group * Day + (1|subject) + (1|stimID).`

Given that the model of hippocampal pattern separation predicts that lower NPS (particularly for semantically similar pairs) (Favila et al., 2016), we expected that group differences in NPS should be associated with worse subsequent memory in hippocampus, reflecting a failure to differentiate overlapping representations. Multiple comparisons were corrected using the Benjamini-Hochberg false discovery rate procedure within each ROI and model.

## Results

### Behavior

#### Memory Performance

In order to investigate the relationship between treatment group and treatment day on performance on the two memory tasks (conceptual and perceptual recognition memory), we used linear mixed-effects models. Our analyses revealed a clear difference between the two task types. In the conceptual memory task, when comparing performance on day 4 (the final treatment day) and day 1 (the baseline day with no treatment), we found a significant interaction between group and day (β = 0.37, z = 2.8, p = 0.0049) in predicting hits (i.e. responding “old” to a stimulus that was presented at encoding). Unexpectedly, a similar result was seen for false alarms (i.e. responding “old” to a stimulus that was not presented at encoding) (β = 0.48, z = 2.15, p = 0.032). In the perceptual memory task, no significant difference was found between treatment groups or treatment days for either hits (β = –0.076, z = –0.49, p = 0.63) or false alarms (β = – 0.27, t = –1.23, p = 0.22).

Together these models show that TMS treatment increased both hits and false alarms in the active group on the conceptual memory task. And yet, neither task showed a significant criterion shift by day 4 (Conceptual: t = 1.97, p = 0.069; Perceptual: t = 1.02, p = 0.33) (**Supplement S2**). Therefore, while TMS treatment appears to increase the likelihood that active-treatment subjects will rate an image as “old,” which increases their rates of hits and false alarms, this effect does not meet the standards for a criterion shift.

#### Effect of treatment on response bias

We next considered the response during retrieval (i.e., “old” vs. “new”) regardless of accuracy. We found that there is a significant group:day interaction for conceptual memory (β = 0.38, z = 3.7, FDR-corrected p-value = 0.0017) but not for perceptual memory (β = 0.14, z = 1.14, FDR-corrected p-value = 0.41) nor for encoding relatedness judgments (β = 0.06, z = 0.22, FDR-corrected p-value = 0.79). Taken all together, these models help contextualize the behavioral results. While TMS treatment does not affect the active and control groups differently in terms of their relatedness judgment at encoding, they show a clear difference in conceptual memory and not perceptual memory. Active subjects are more likely to say “old” than the control group by the end of treatment, which helps explain the observed increase in both hits and false alarms.

### Relationship between treatment group, treatment course, and representational strength on successful memory

#### Effect of TMS Treatment on Semantic RSA

We first assessed the effect of TMS on the active treatment participants, comparing the pre-treatment baseline with the post-treatment scan after three consecutive daily TMS sessions. We used the LLM Llama-3.2-3b layer 6 to generate the semantic model RSM and VGG16 Layer 3 to generate the visual model RSM. We then found the correlations between these RSMs to neural RSMS to determine semantic and visual representational strength. It was immediately clear that in inferior parietal lobule, TMS increased semantic representational strength values relative to baseline but has no apparent effect on visual representational strength (**Figure 5**). We then employed linear mixed-effects models to investigate the relationship between treatment group (active vs control) and treatment course (baseline day vs post-treatment) on semantic representational strength. For semantic RSA, we found that both ROIs showed a significant group:day interaction after Bonferroni correction: inferior parietal lobule (t = 2.38, BH-adjusted p = 0.034, 95% CI: [0.0024, 0.025]) and hippocampus (t = –2.46, BH-adjusted p = 0.027, 95% CI: [–0.027, –0.003]. On the other hand, for visual RSA we found that neither ROI showed a significant group:day interaction after Bonferroni correction: inferior parietal lobule (t = –0.01, BH-adjusted p = 0.88, 95% CI: [–0.014, 0.012]) and hippocampus (t = 0.022, BH-adjusted p = 0.9, 95% CI: [–0.018, 0.032]). Interestingly, this shows that the TMS treatment resulted in increased semantic representational strength at the parietal stimulation site and decreased semantic representational strength in the downstream MTL target (**Figure 5**).

**Figure 5.**
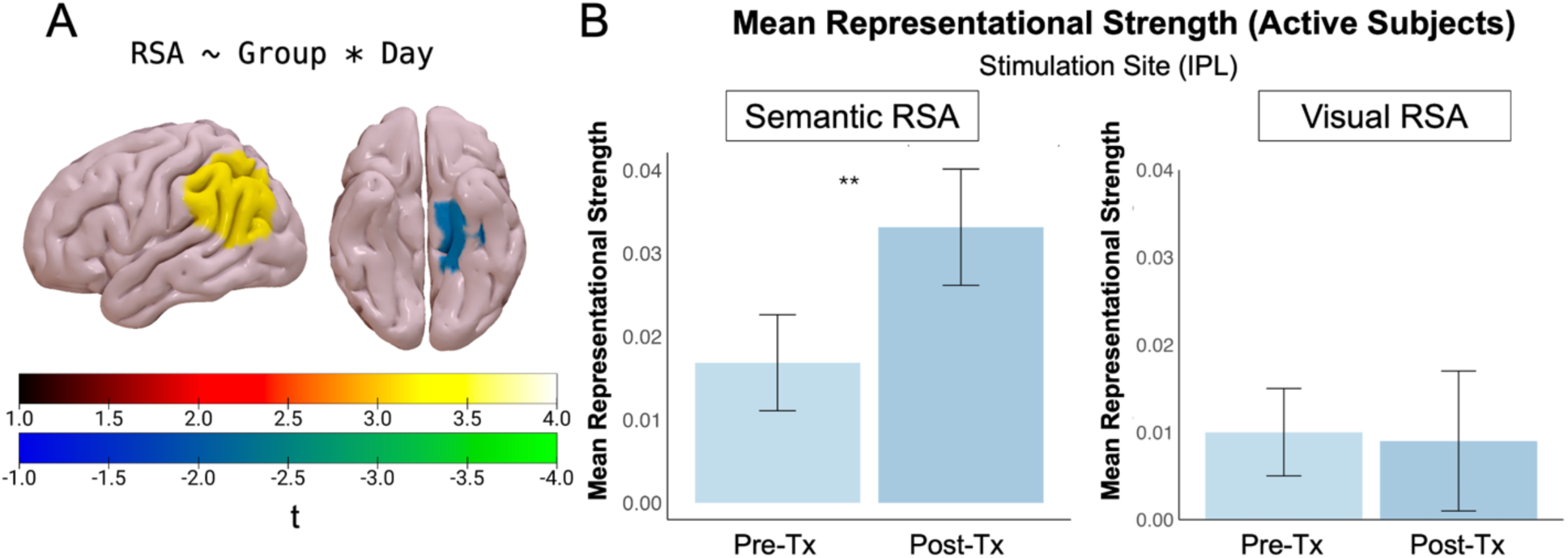
Effect of TMS on Representational Strength. (A) Two-way interaction between treatment group (active vs control) and treatment course (pre-treatment vs post-treatment) for all trials regardless of subsequent memory. (B) For all active stimulation participants, mean representational strength at Inferior Parietal Lobule stimulation site generated with a model RSM based on Llama-3.2-3b layer 6 with Euclidean Distance for semantic RSA and a model RSM based on VGG16 Layer 3 with Euclidean Distance for visual RSA.

#### Relationship between TMS treatment and memory performance

To test whether TMS enhances semantic representations during encoding in a manner that supports successful memory, we next examined Group:Day interactions for subsequently remembered trials only. (We discuss the relative merits of such an approach compared with a Group:Day:Memory interaction model in **Methods** and present three-way interaction results in **Supplement 3**). We found memory-related increases in semantic representational strength in the active TMS group from pre- to post-treatment at the stimulation ROI inferior parietal lobule (t = 2.61, BH-adjusted p = 0.018, 95% CI: [0.0045, 0.031]). In Hippocampus, on the other hand, we found memory-related decreases in semantic representational strength in the active group over treatment for subsequently remembered trials (t = –2.38, BH-adjusted p = 0.034, 95% CI: [–0.032, –0.003]). Therefore, it appears that the effect of TMS on memory-related semantic representational strength increased in some regions but not others (**Figure 6**).

**Figure 6.**
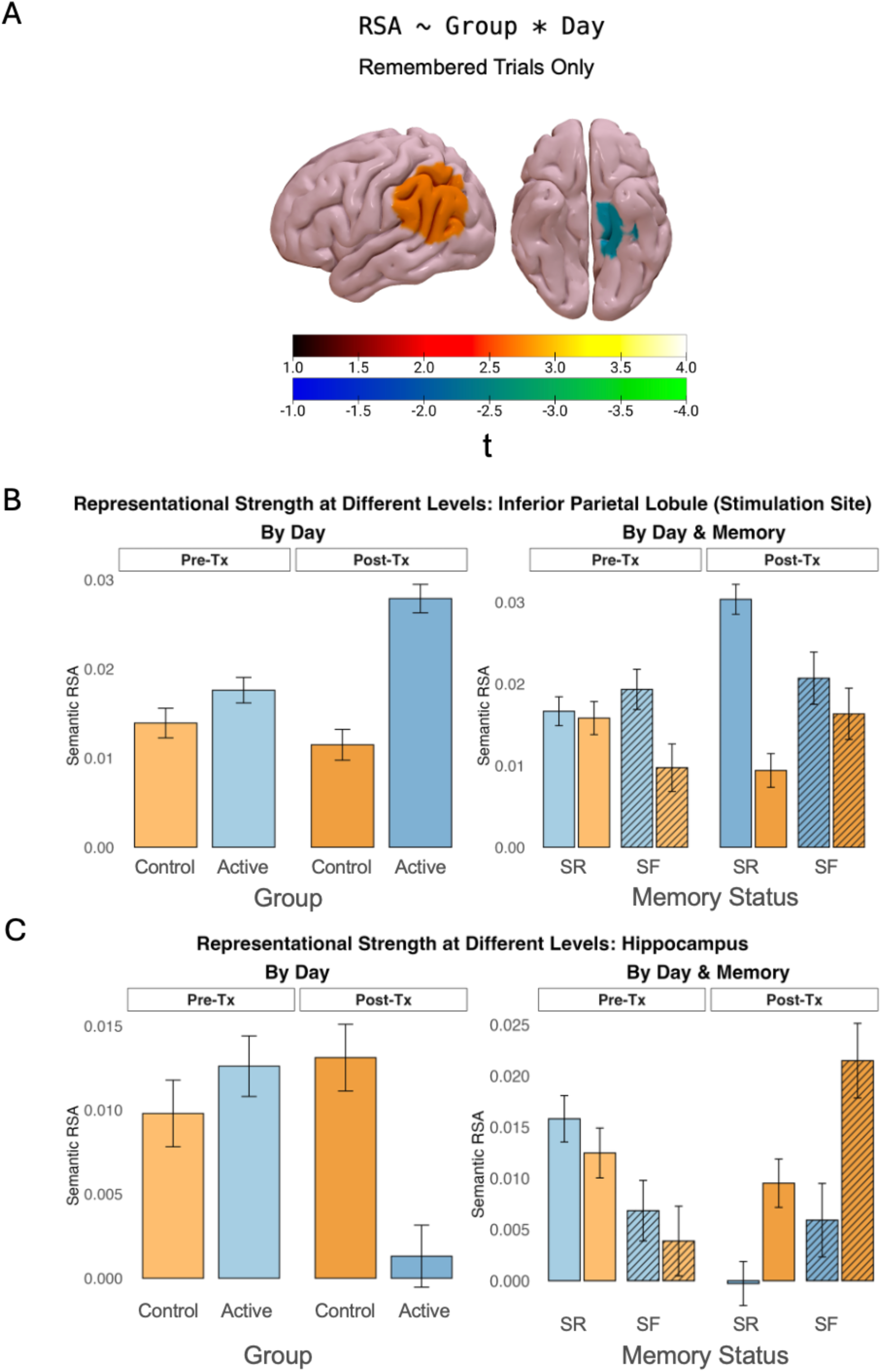
Relationship between Group, day, Representational Strength, and Memory Performance. (A) Two-way interaction between treatment group (active vs control) and treatment course (baseline/day 1 vs post-treatment/day 4) for subsequently remembered trials only. (B) Overall semantic RSA results split by task conditions in Inferior Parietal Lobule for active (n=10) and control (n=9) subjects split by treatment day and group (left) and treatment day, group, and subsequently remembered (SR) and subsequently forgotten (SF) trials (right). (C) Same condition split for Hippocampus.

Taken together, we can see that TMS increases semantic representational strength in parietal cortex, perhaps supporting the integration of semantic information into memory. In hippocampus, there is a clear functional difference. We found that reduced representational strength supports successful memory (**Figure 7**). These findings may indicate a difference in object-level semantic processing between the two regions. Such a pattern is consistent with previous work that has shown that pattern separation and differentiation are required for successful memory (Favila et al., 2016; Karlsson Wirebring et al., 2015) and we will explore this concept further in the following section.

**Figure 7.**
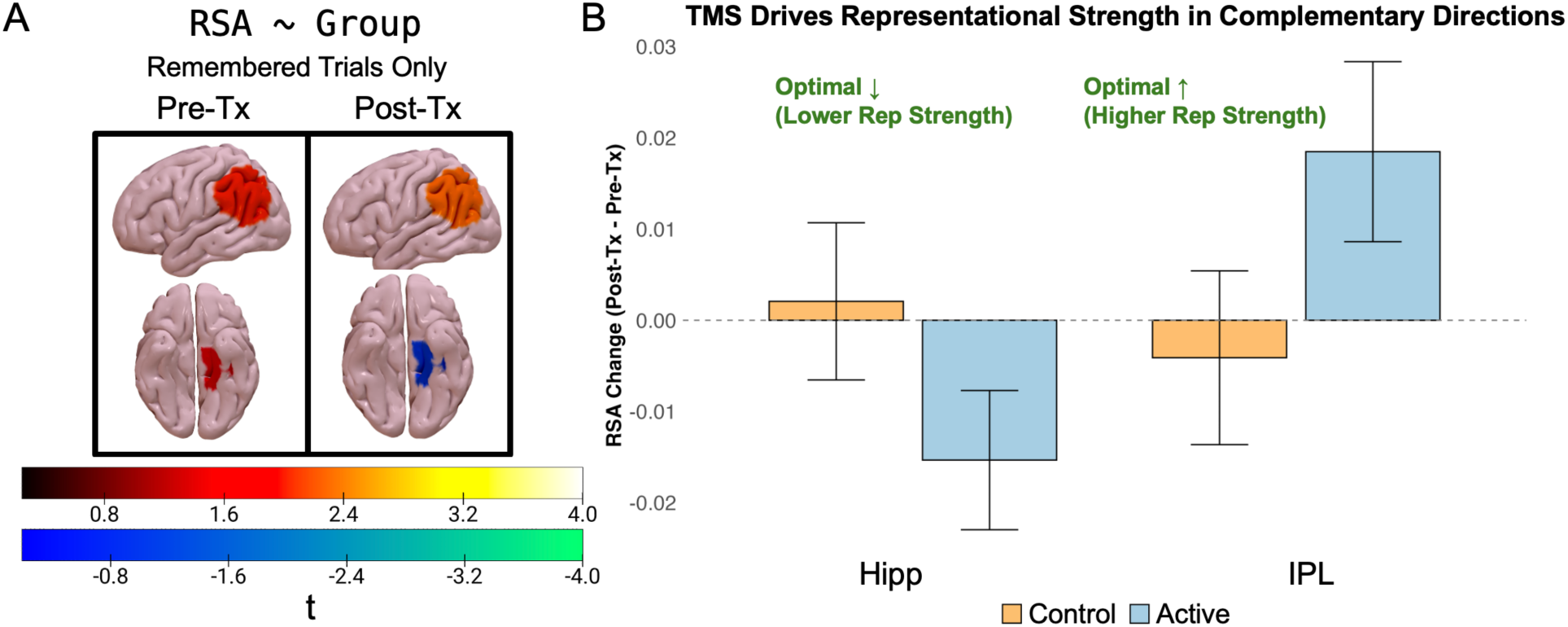
Summary of treatment effects on RSA results in the four regions with significant memory effects. (A) ROIs showing t-values that reflect the slope of the relationship between group and representational strength for remembered trials only, pre- and post-treatment (B) Difference between representational strength post-treatment vs pre-treatment. Increased representational strength in IPL promotes better memory, whereas decreased representational strength in hippocampus promotes better memory.

### Relationship between neural pattern similarity, RSA, and memory

So far, we have shown that less representational strength in hippocampus promotes better memory and hypothesized that this reduced representational strength could be capturing pattern separation mechanisms that are known to be important for successful memory. And yet, we do not have sufficient evidence to support this claim because of a feature of RSA. RSA is a second-order correlation between similarity in a model RSM or neural RSM (or dissimilarity in RDMs) and cannot disambiguate first-order correlations within the model matrices. Thus, a finding of “low representational strength” could indicate low neural similarity that fails to correlate with the similarity in the model, or low similarity failing to index a high level of neural similarity.

Therefore, RSA is not as well-suited to assess pattern separation in hippocampal activity as other methods, such as neural pattern similarity (NPS). This measure captures the similarity of dissimilarity between voxel-wise activity within each ROI, and thus higher or lower NPS (i.e. cortical distinctiveness, c.f. Yu et al., 2024) is a more direct measure of the presence or absence of pattern separation.

We adapted a method used by Wing et al. (Wing et al., 2020) to investigate the relationship between NPS within ROIs and treatment group and day on memory success. We calculated the mean correlation between the single-trial beta for each item and the items from different encoding runs and used linear mixed-effects models to investigate the interaction between NPS, group, and day. First, we looked at the main effect of NPS of memory and found a significant effect in IPL (z = 2.78, p = 0.006) but not in hippocampus. Greater NPS at the stimulation site appears to promote successful memory. We then looked at the interaction of NPS and Group and found two significant effects in IPL (z = –2.78, p = 0.005) and Hippocampus (z = 2.86, p = 0.004) (**Figure 8**).

**Figure 8.**
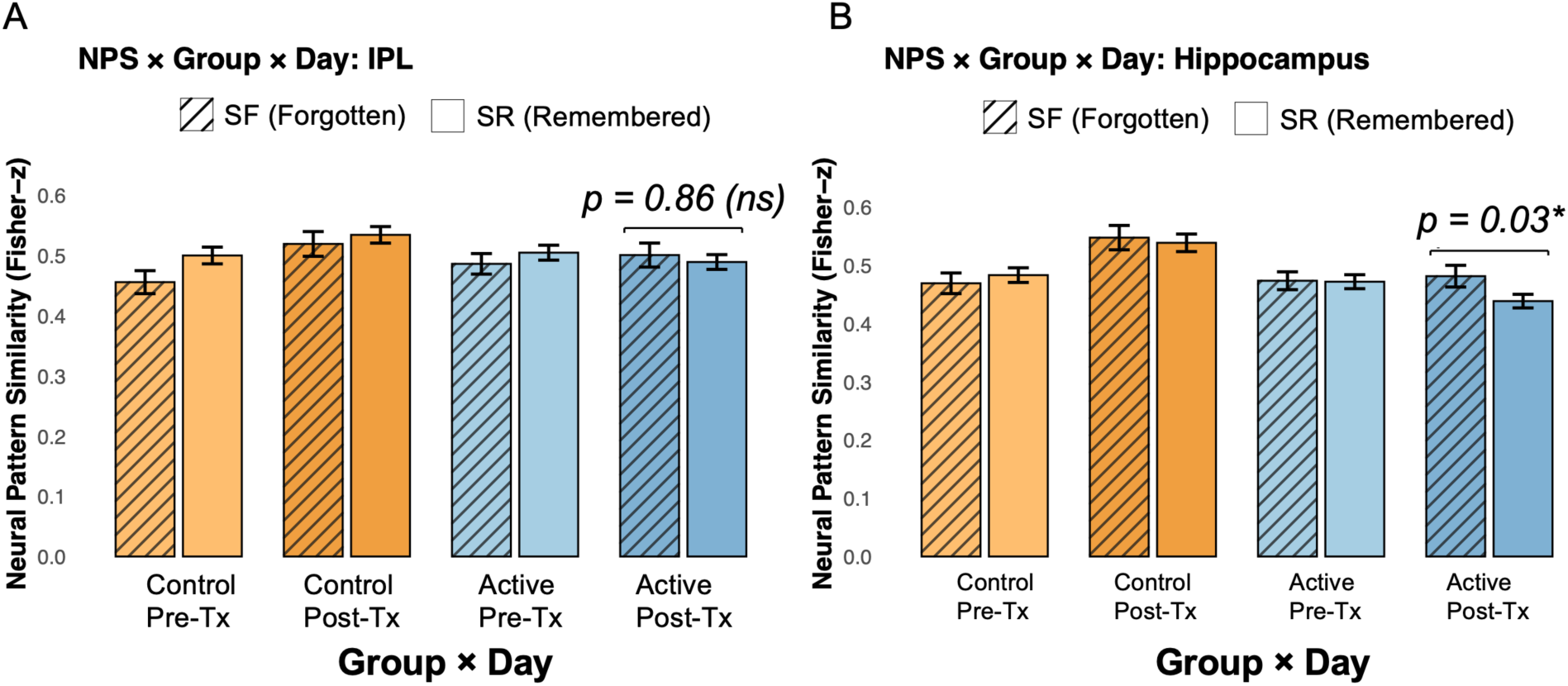
Neural Pattern Similarity (NPS) between each item and all other items in other encoding runs, split by group, treatment day, and subsequent memory at conceptual retrieval. (A) Inferior Parietal Lobule, no significant difference between remembered and forgotten trials in the active group post-treatment. (B) Hippocampus, the active group post-treatment shows significantly lower NPS for remembered trials than for forgotten.

Interpreting the signs and magnitudes of these interaction slopes, we can see that while in IPL, the relationship between NPS and memory is weaker for the active group (i.e. they are less dependent on neural similarity for memory success), in hippocampus we see the opposite. The relationship between NPS and memory is greater in the active group and are therefore more dependent on neural similarity for successful memory. While the interaction models that included day were not significant, we can see that by the final treatment day, the active group shows less NPS for subsequently remembered items than for subsequently forgotten and there is no such different pre-treatment. This provides some support for the claim that neural similarity must be reduced in hippocampus for successful memory, though future work will need to explore this relationship further.

## Discussion

We have presented evidence that TMS affects memory performance and semantic representations in older adults with Mild Cognitive Impairment. These results corroborate previous research showing that the mnemonic benefits of iTBS involve functional changes in parietal and hippocampal/parahippocampal regions (Hermiller, 2024; Hermiller et al., 2019) and extend this understanding by demonstrating that semantic information processing has region-specific effects on recognition memory. In particular, we present evidence that parietal TMS differentially affects the stimulation site and downstream hippocampal target in a way that supports successful memory.

We found that both hit rate and false alarm rate were affected by TMS treatment, where the active treatment group shows an increase in both hits and false alarms over the course of treatment compared to the control group. Moreover, we showed clear evidence that the active stimulation group shows a bias toward rating items as old (i.e. increasing both hits and false alarms) relative to control. The increase in false alarms was unexpected and suggests that such a change could be an unintentional effect of TMS treatment. If TMS is indeed affecting the availability and/or processing of semantic information, it is reasonable to infer that it can lead to an increased sense of familiarity for the objects among active treatment participants relative to the control participants. This would explain why participants tended to rate items as old more often, leading to increased rates of both hits and false alarms.

We next observed that TMS treatment affects semantic representational strength, leading to a post-treatment increase in inferior parietal lobule (which contains AG) and post-treatment decrease in hippocampus. When we considered the relationship between the changes in semantic representational strength and memory performance, we saw that the increase in parietal representational strength and decrease in hippocampal representational strength were associated with better memory. The increase in parietal representational strength is not surprising given what is known about the role of lateral parietal cortex in memory formation (Dave et al., 2022). It has been previously shown that increased activity in lateral parietal cortex is associated with successful memory encoding (de Chastelaine et al., 2016) and retrieval (Howard et al., 2024; Rugg & King, 2018; Yazar et al., 2014) due to its role in integrating information from multiple sensory modalities (Shimamura, 2011). This abstracted information is thought to draw from sources of experiential information, coming from sensory, motor, spatial, or affective modalities (Binder et al., 2016; Fernandino et al., 2022). As such, one would expect an increase in semantic representational strength in AG to be associated with improved memory, as greater fidelity of stimulus information would be available for encoding.

We also found significant modulation of representations in parahippocampal and hippocampal cortices, regions that are distant from the parietal stimulation site, but nonetheless play a critical role in memory encoding. Previous work has shown that TMS applied to dorsolateral prefrontal cortex increases stimulus representation in anterior temporal lobe and hippocampus in healthy participants (W.-C. Wang et al., 2018), and the current work extends this phenomenon to lateral parietal neuromodulation. The reduced representational strength in the hippocampus we observed is suggestive of pattern separation, a critical function of the hippocampus (Favila et al., 2016; Leal & Yassa, 2018). We then found that lower neural pattern similarity (i.e. increased distinctiveness) is associated with improved memory in active subjects post-treatment. This lends support to the idea that hippocampus must limit similarity between item representations in order to support successful recognition, though this is only a preliminary step toward a more complete account of the effect of TMS on semantic representations in hippocampus. Given that TMS increases representational strength in parietal cortex and decreases representational strength in hippocampus regardless of memory and that such changes are related to successful memory, future work is needed to determine how and why TMS treatment affects these regions differently.

We commented above that the effect of TMS on false alarms could be due to an increased sense of familiarity for the objects leading to a change in response bias toward rating objects as “old.” It is well known that memory can be improved through deep semantic processing (Craik & Lockhart, 1972) or by giving instructions to attend to relationships between stimuli (Hunt & Einstein, 1981). When similarity between targets and lures is too high, however, relatedness can lead to false memory (McDermott & Roediger, 1998; Roediger & McDermott, 1995). Such a dynamic, where stimulus information can improve memory up to a point beyond which it interferes with memory may be involved in the false alarm effect we see in our study. While we expected that stimulating AG would improve memory by increasing the availability of semantic information related to the stimuli, this relationship may not be entirely unidirectional.

To that end, there are still many open questions about the role of AG in memory (Humphreys et al., 2021; Humphreys & Tibon, 2023). Several studies have shown how AG, along with parahippocampal cortex, supports recognition memory (Vilberg & Rugg, 2009; Yonelinas et al., 2005), though activity in posterior parietal cortex has been shown to be greater for hits and false alarms than for misses and correct rejections (Wagner et al., 2005). Furthermore, activity in parietal regions (e.g. inferior parietal lobule and precuneus) and medial temporal lobe regions (e.g. hippocampus) differentiates true memory from false memory, though these relationships can depend on confidence as well as accuracy (Urgolites et al., 2015). Other studies have shown that retrieval success, precision, and vividness are experimentally dissociable (Richter et al., 2016) and associated with greater activity in AG (Kuhl & Chun, 2014). It has also been shown that TMS applied to AG can differentially affect memory accuracy and confidence (Grob et al., 2024; Yazar et al., 2014). Therefore, while it is reasonable that an increase in the availability of semantic information at memory encoding would lead to improved recognition memory performance, it may be that such increased semantic information could be affecting other aspects of memory, such as a sense of familiarity, which can interfere with successful recognition.

## Conclusion

In summary, we have presented evidence that parietal TMS affects semantic representations in such a way that supports successful recognition memory in a clinical population. Our account extends previous work by adding further detail about the role of stimulus representations in mnemonic neuromodulation. Based on the role of the angular gyrus as a semantic hub, we suggest that the application of iTBS increased the capacity of lateral parietal cortex to process the representation of semantic information, leading to improved memory performance.

Furthermore, the effect of parietal stimulation on hippocampus is to *reduce* stimulus representations, perhaps to promote pattern separation, which is in turn necessary for successful memory. Future work is needed to expand our understanding of how TMS affects the encoding of stimulus information and its impact on memory in order to inform future clinical interventions.

## Supplement

### Model RSM Distance Metric Selection

While correlation-based metrics (Pearson, cosine) are standard in RSA, they don’t capture magnitude information, (Kriegeskorte & Kievit, 2013). While this limitations can be addressed with data preprocessing (Clarke et al., 2024) or robustness checks with additional distance metrics (Jiang et al., 2024), in other cases, researchers may select Euclidean or Mahalanobis distance (Levy et al., 2024), or use a combination of models depending on what is required to capture the representational geometry of the neural activity space (Khaligh-Razavi & Kriegeskorte, 2014).

Given that LLM embeddings encode semantic relationships through both direction and magnitude in high-dimensional space (3072 dimensions), we evaluated which distance metric maximized signal-to-noise ratio (SNR = between-subject variance / within-subject variance) for a random subset of our neural RSA data (i.e. the trial RSA values for the IPL and Hippocampus ROIs). Using the sample() function in R [cite] we held out 10% of all trials across subjects for distance metric selection. Trial-level sampling was chosen to include inter-subject variability in the analysis.

Among eight candidate metrics (Pearson correlation, normalized correlation, cosine similarity, Spearman correlation, Euclidean, Manhattan, Minkowski, dot product), the Euclidean-family distance metrics (Euclidean, Manhattan, Minkowski) showed substantially higher SNR than correlation-based metrics (Figure S1). Though it is not yet known which method is most suitable for neural data, we selected Euclidean distance (SNR=0.0132) based on theoretical work that suggests that Minkowski distance (though performing similarly with an SNR of 0.0132) may not be as effective for high-dimensional data compared with Euclidian distance, depending on the implementation (France et al., 2012). For our purposes, we suspect that magnitude-preservation in all three of our top-performing models (as indexed by SNR) may better capture the high-dimensional embedding space of the model RSMs and perhaps the semantics-neural relationship RSA is meant to measure.

**Figure S1.**
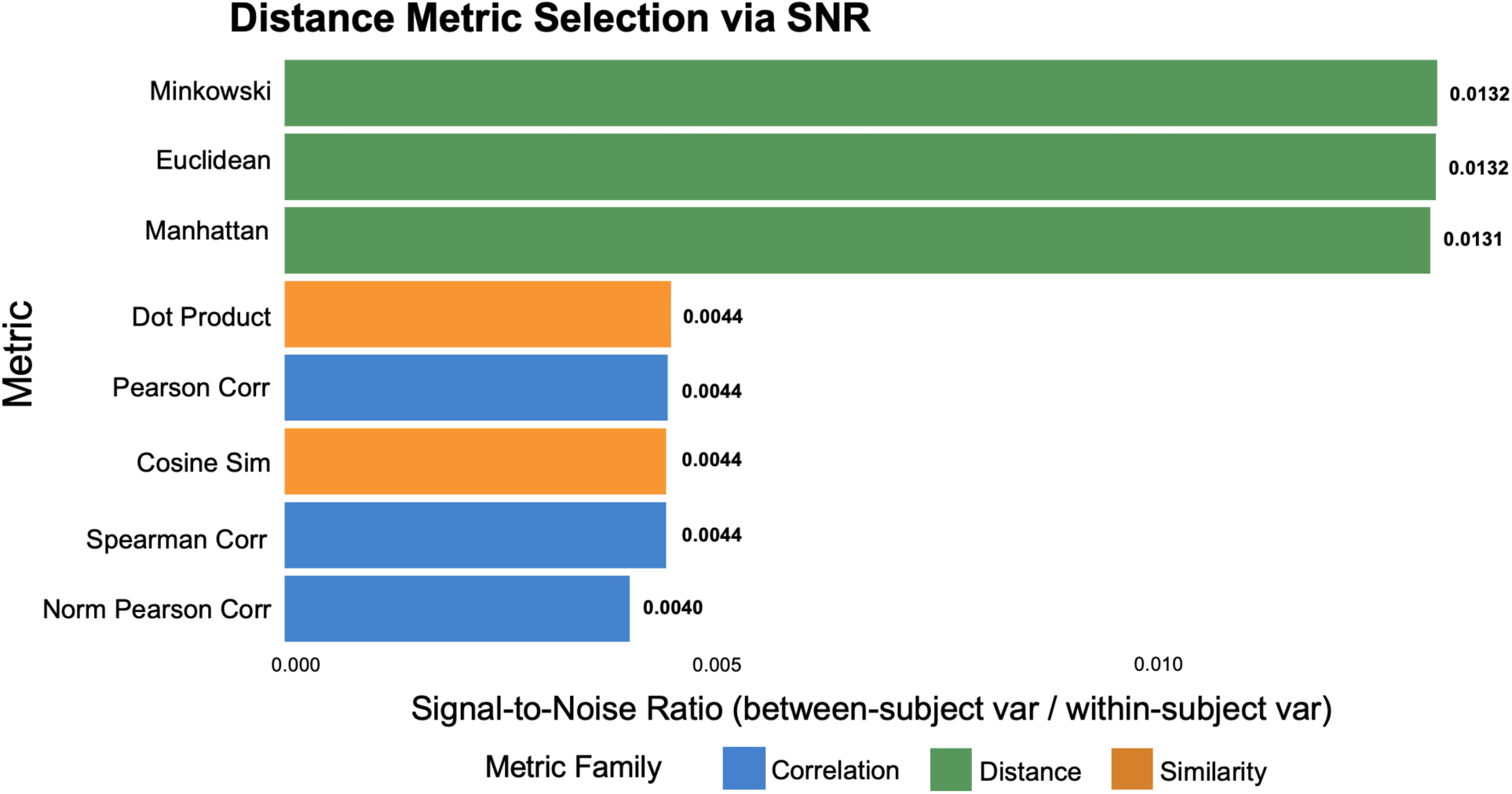
(A) Signal-to-Noise Ratio (SNR) (`between-subject variance / within-subject variance`) for each distance metric across entire data set. (B) Stability analysis showing relative SNRs for each distance metrics with progressively larger subsamples

### Supplemental Figure 2. Criterion Analysis

**Figure S2.**
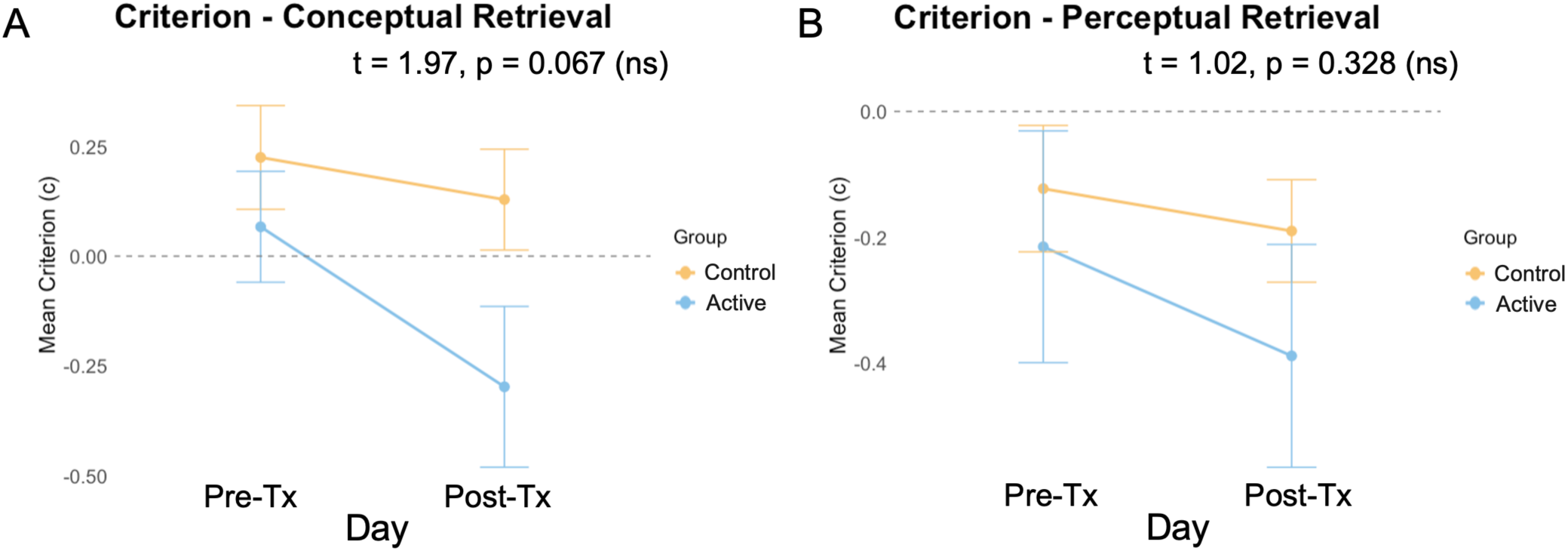
Response bias (criterion) defined as `–0.5× [z(HR) + z(FAR)]`. (A) Criterion pre- and post-treatment for conceptual retrieval. (B) Criterion pre- and post-treatment for perceptual retrieval

### Supplement 3: Three-way interaction model results

We discussed in the Methods section why we find two-way Group:Day models for subsequently remembered trials preferable to three-way Group:Day:Memory interaction models because a three-way interaction would tell us the relationship between the Group:Day slope and memory, which answers a related but distinct question to the effect of treatment group and day on memory, while maintaining representational strength as the (continuous) outcome variable. We ran three-way models as well and found that in IPL, results are similar but less strong (t = 2.11, BH-adjusted p = 0.049, 95% CI: [0.002, 0.054). For Hippocampus we see a similar result where the results are not significant (t = –1.54, BH-adjusted p = 0.023, 95% CI: [–0.056, 0.0067]). While one could argue that because the three-way interaction models are less strong (i.e. one is barely significant and the other is not), our results are less meaningful, we attribute this difference to the comparatively less sensitive nature of the three-way interaction model in answering our question. Still, it is important that these results should give us some skepticism in interpreting the two-way interaction models, assuming that a stronger effect would be significant in both cases.

